## SupplementaryInformation for "Engineering a symbiont as a biosensor for the honey bee gut environment"

---

### Supplementary Figures

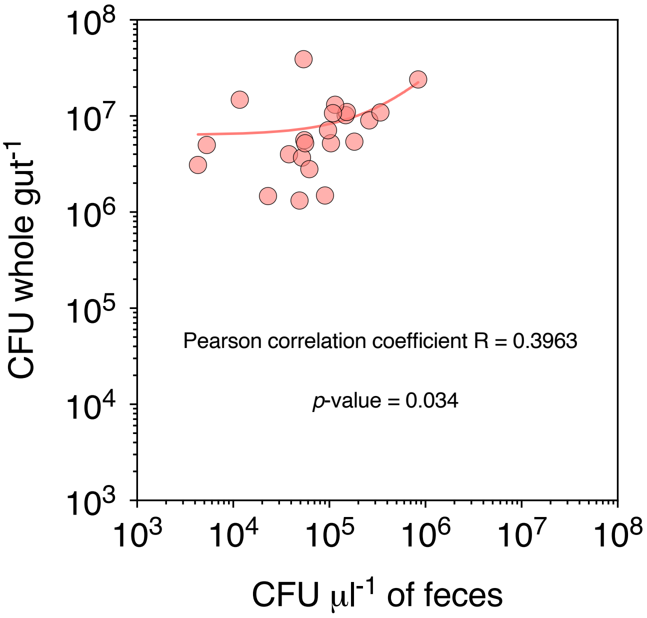


**Supplementary Figure 1. Bacterial load from feces is a proxy for levels of gut colonization.** Scatterplot shows linear regression of bacterial concentration values of engineered *S. alvi* found in matching samples of feces and gut homogenates (*i.e.* feces and gut were sourced from the same bee). Pearson correlation coefficient R and *p*-value are provided, for $n = 22$.

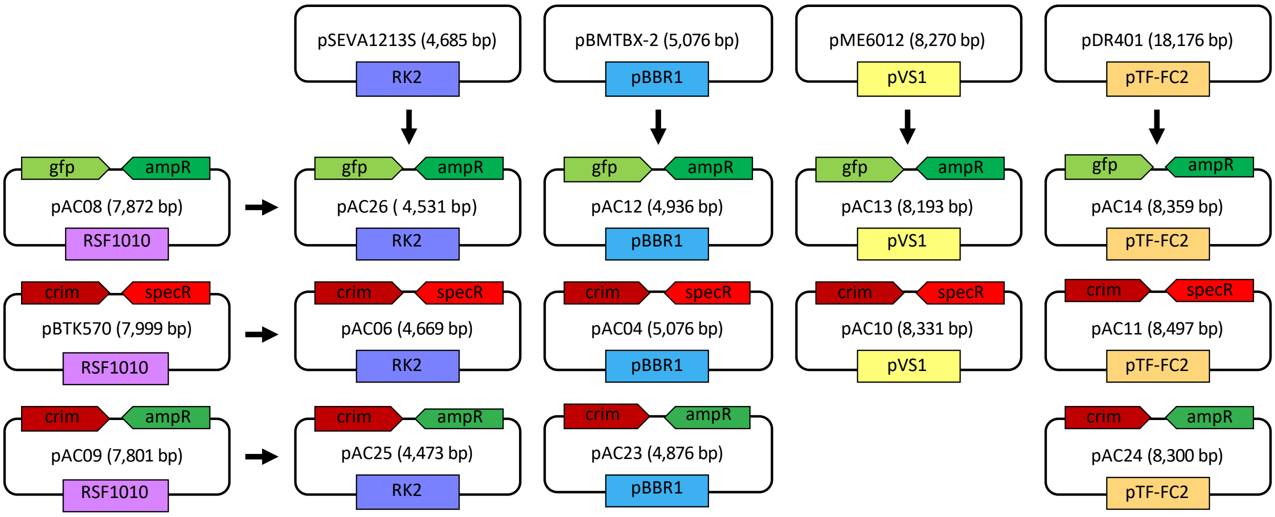


**Supplementary Figure 2. Plasmid maps of the broad-host range vectors developped in this study.** Broad-host-range replicons were sourced from the pSEVA1213S, pBMTBX-2, pME6012 and pDR401 plasmids (top row). The standard fragments bearing the antibiotic marker and fluorescent protein were obtained from the RSF1010-based vectors pBTK570 (Leonard *et al.*, 2018), pAC08 and pAC09 (left column). In those, RSF1010 was replaced with the different broad-host range replicons, resulting in the eleven new vectors shown.

**Supplementary Table 1**. **Plasmids used in this study.**

| Plasmid | Description | Reference or source |
|---|---|---|
| pBTK503 | pBTK plasmid with constitutive GFP, RSF1010 replicon | Addgene No. 110616 [27] |
| pBTK570 | pBTK plasmid with constitutive E2-crimson, RSF1010 replicon | Addgene No. 110615 [27] |
| pBTK552 | pBTK plasmid with IPTG-inducible GFP, RSF1010 replicon | Addgene No. 110618 [27] |
| pBMTBX-2 | Template plasmid carrying the pBBR1 replicon | Addgene No. 26073 [33] |
| pME6012 | Template plasmid carrying the pVS1 replicon | Heeb *et al*., 2000 [35] |
| pDR401 | Template plasmid carrying the pTF-FC2 replicon | Rawlings *et al.*, 1984 [37] |
| pSEVA1213S | Template plasmid carrying the RK2 replicon | Addgene No. 122095 [31] |
| pAC08 | Constitutive GFP, ampR, RSF1010 replicon | This study |
| pAC09 | Constitutive E2-crimson, ampR, RSF1010 replicon | This study |
| pAC26 | Constitutive GFP, ampR, RK2 replicon | This study |
| pAC06 | Constitutive E2-crimson, specR, RK2 replicon | This study |
| pAC25 | Constitutive E2-crimson, ampR, RK2 replicon | This study |
| pAC12 | Constitutive GFP, ampR, pBBR1 replicon | This study |
| pAC04 | Constitutive E2-crimson, specR, pBBR1 replicon | This study |
| pAC23 | Constitutive E2-crimson, ampR, pBBR1 replicon | This study |
| pAC13 | Constitutive GFP, ampR, pVS1 replicon | This study |
| pAC10 | Constitutive E2-crimson, specR, pVS1 replicon | This study |
| pAC14 | Constitutive GFP, ampR, pTFC-FC2 replicon | This study |
| pAC11 | Constitutive E2-crimson, specR, pTFC-FC2 replicon | This study |
| pAC24 | Constitutive E2-crimson, ampR, pTFC-FC2 replicon | This study |
| pAC17V5a | IPTG-inducible two-plasmid system bearing CP25 > lacO-GFP, specR, RSF1010 replicon (1/2) | This study |
| pAC17V5b | IPTG-inducible two-plasmid system bearing CP25 > lacI, ampR, pTF-FC2 replicon (2/2) | This study |

**Supplementary Table 2. Primers used in this study.** Uppercase letters indicate priming sequence and lowercase nucleotides show homology regions for Gibson assembly as primer overhangs.

| Primers | Sequences (5' – 3') | Description | Resulting plasmid |
|---|---|---|---|
| AC09 | ATCTGAATCATGCGCGGATG | Linearize pAC08/pAC09/pBTK570 without replicon | N/A |
| AC10 | CACGTTAAGGGATTTTGGTCATG | | |
| AC16 | gaccaaaatcccttaacgtgCCAAAGGGTTCGTGTAGACT | Amplify RK2 from pSEVA1213S | pAC06 / pAC26 / pAC25 |
| AC17 | catccgcgcatgattcagatGCGCGCCGTAGAAAAGATC | | |
| AC11 | gaccaaaatcccttaacgtgACGCCCGGTAGTGATCTTAT | Amplify pBBR1 from pBMTBX-2 | pAC12 / pAC04 / pAC23 |
| AC12 | catccgcgcatgattcagatGACGAGCCTCAGACTCCAG | | |
| AC48 | catccgcgcatgattcagatCTGGTTTCGGAGTTGACAGC | Amplify pTF-FC2 (fragment 1/2) from pDR401 | pAC14 / pAC11 / pAC24 / pAC17V5b |
| AC49 | CGTCTTTGATAAGCCGCTCC | | |
| AC50 | GGAGCGGCTTATCAAAGACG | Amplify pTF-FC2 (fragment 2/2) from pDR401 | |
| AC51 | gaccaaaatcccttaacgtgCTCAAGGCAGCCAGAACATC | | |
| AC46 | GAAACCGCCATCAGTACCAG | Amplify pVS1 (fragment 1/2) from pME6012 | pAC13 / pAC10 |
| AC44 | gaccaaaatcccttaacgtgTTCCTGGCGTTTTCTTGTCG | | |
| AC45 | catccgcgcatgattcagatGTAGACAACATCCCCTCCCC | Amplify pVS1 (fragment 2/2) from pME6012 | |
| AC47 | CTGGTACTGATGGCGGTTTC | | |
| AC36 | CGAGCGGCCGCGATTATC | Linearize pBTK503 (fragment 1/2 for pAC08 and 1/3 for pAC09) | pAC08 / pAC09 |
| AC39 | GTCAAGCGCTTCGGTCATG | | |
| AC37 | ttgataatcgcggccgctcgCCACTGACGAGCAGATTTCCAG | Linearize pBTK503 (fragment 2/2) for pAC08 | pAC08 |
| AC38 | CATGACCGAAGCGCTTGAC | | |
| AC41 | ttgataatcgcggccgctcgAGTATCCGGTTCCCATCTGC | Linearize pBTK570 (fragment 2/3) | pAC09 |
| AC42 | ctttggcagtttattcttgacatgtagtgagggggctggtataatcacatagtactgtt ATACAGAAACAGAGGAGATATTACATATGG | | |
| AC43 | ttgataatcgcggccgctcgCCACTGACGAGCAGATTTCCAG | Linearize pBTK570 (fragment 3/3), to use with AC38 | |
| AC_59 | GGTAGCCATTGATTGCCTCC | Linearize pBTK552 (fragment 1/2) | pAC17V5a |
| AC_114 | gtagtcggcaaataagcggcGTGAACACTCTCCCGGTTGT | | |
| AC_60 | GGAGGCAATCAATGGCTACC | Linearize pBTK552 (fragment 2/2) | |
| AC_70 | GCCGCTTATTTGCCGACTAC | | |
| AC_115 | gatgacacgaactcacgacgCGCTCGGCTTTGGCAGTTTATTC | Amplify *lacI* from pBTK552, to clone with pTF-FC2 | pAC17V5b |
| AC_116 | ttgataatcgcggccgctcgAAACAAAAGGCCCAGTCTTCC | | |
| **Primers** | **Sequences (5' – 3')** | **Description** | **Reference or source** |
| AC30 | gtgggaatctacctatttctacg | qPCR primers binding the 16S region of *B. apis* | Kesnerova *et al.*, 2017[51] |
| AC31 | aacgcgggctcatctatctc | | |
| AC28 | cttagagataggagagtgcctt | qPCR primers binding the 16S region of *S. alvi* | Kesnerova *et al.*, 2017[51] |
| AC29 | aacttaatgatggcaactaatgacaa | | |
| AC24 | ctccaaggcgtacatcaagc | qPCR primers binding the E2-crimson | This study |
| AC25 | gtcctcgaagttcatcacgc | | |

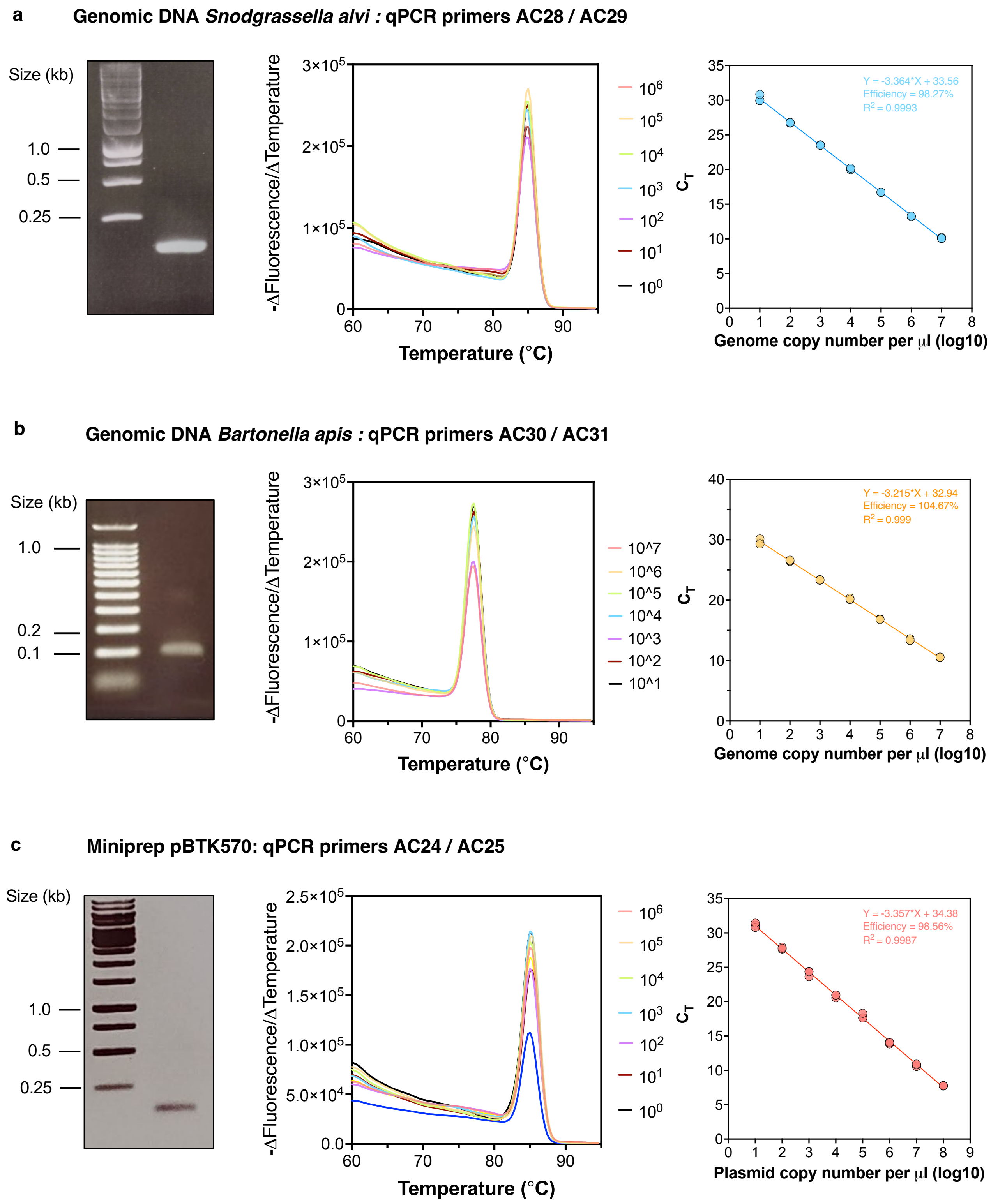


**Supplementary Figure 3. Validation of qPCR primers and standard curves.** Primers specificity was confirmed by visualization of amplicons on an agarose gel (left panel) and generation of melting curves (middle panel). Standard curves were generated using serially diluted genomic DNA of **a** S. alvi, **b** B. apis or **c** miniprep of pBTK570.

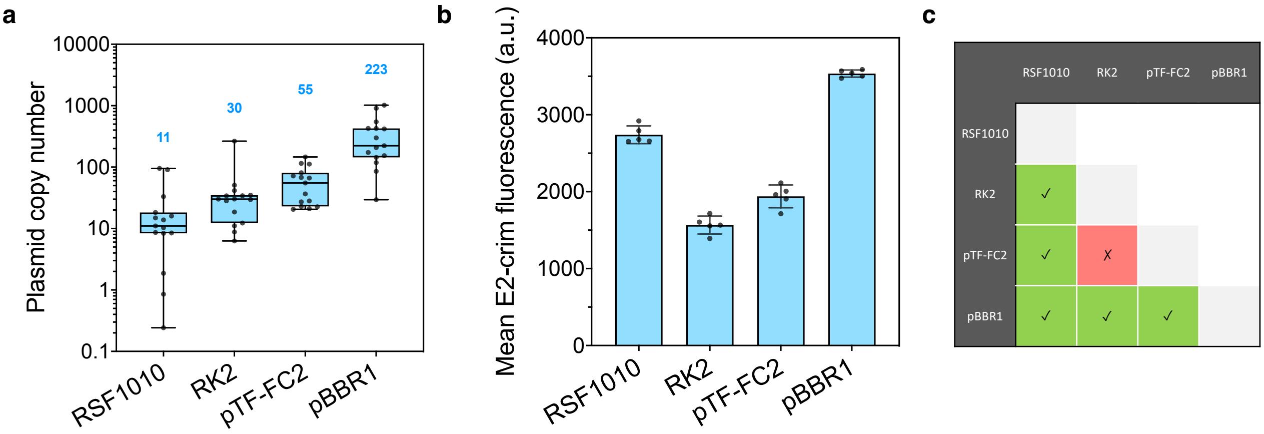


**Supplementary Figure 4. Characterization of functional broad-host range replicons in the honey bee gut symbiont *B. apis.*** **a** Broad-host range plasmids have different copy numbers in *B. apis.* Box plots show median values of plasmid copy numbers obtained by qPCR from three independent experiments with five biological replicate each (total n=15). Median copy number are indicated with the corresponding box plots. **b** The difference in plasmid copy number results in different protein expression levels in *B. apis.* Graph shows mean of E2-crimson fluorescence $\pm$ standard deviations of five biological replicates. Each replicate represents the average fluorescence of at least 9,000 cells measured by flow cytometry. Plasmids used for panels **a** and **b** in *B. apis* were pBTK570, pAC06, pAC11, and pAC04, carrying the RSF1010, RK2, pTF-FC2 and pBBR1 origins of replication, respectively. **c** Some replicons are compatible and can be co-transformed in *B. apis.* Matrix table indicates compatible (green boxes with check mark) and incompatible (red boxes with cross mark) replicons. Vectors were found compatible upon their successful co-transformation by conjugation in *B. apis* cells.

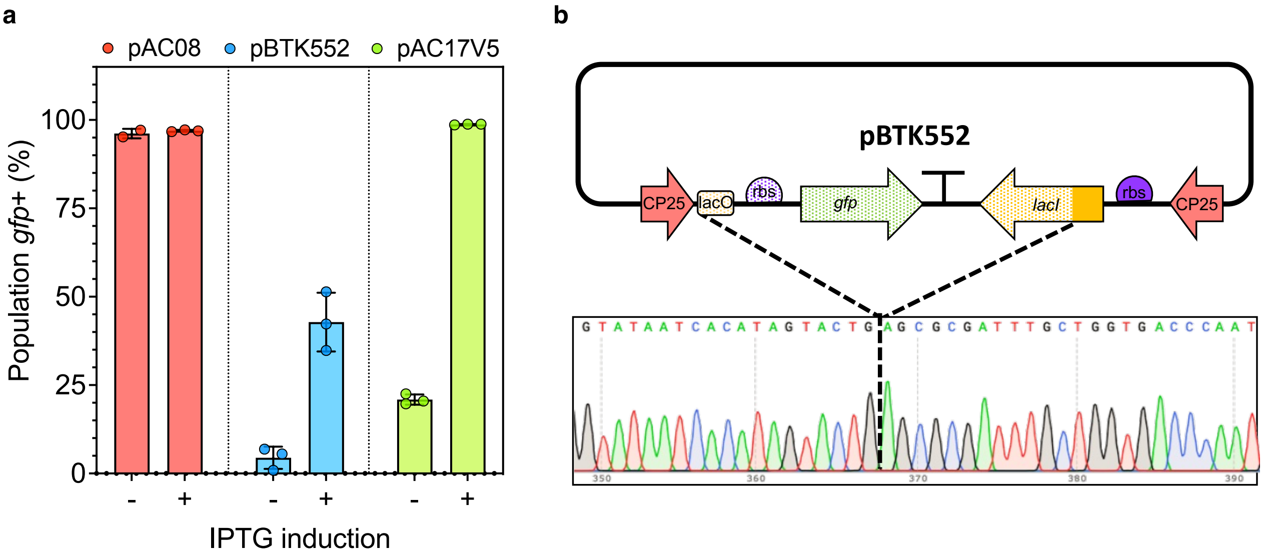


**Supplementary Figure 5. The plasmid pBTK552 (Leonard *et al*., 2018)[27] is genetically unstable in our experimental conditions. a** Graph shows mean ± standard deviation percentage of bacterial population GFP positive. Three biological replicates were tested for each construct. Each replicate value is based on the average fluorescence of at least 9,000 *S. alvi* cells measured by flow cytometry, which were grown in liquid with (+) or without (-) IPTG. As a reference, *S. alvi* bearing the pAC08 plasmid constitutively expressing GFP and *S. alvi* carrying our pAC17V5 dual-vector system were also analyzed. **b** The CP25 inter region gets deleted from pBTK552. Plasmid map of the pBTK552 vector (top panel) with a representative sanger sequencing of a frequent deletion obtained in the unstable region (bottom panel) are shown. Dotted lines indicate the position of the deletion.

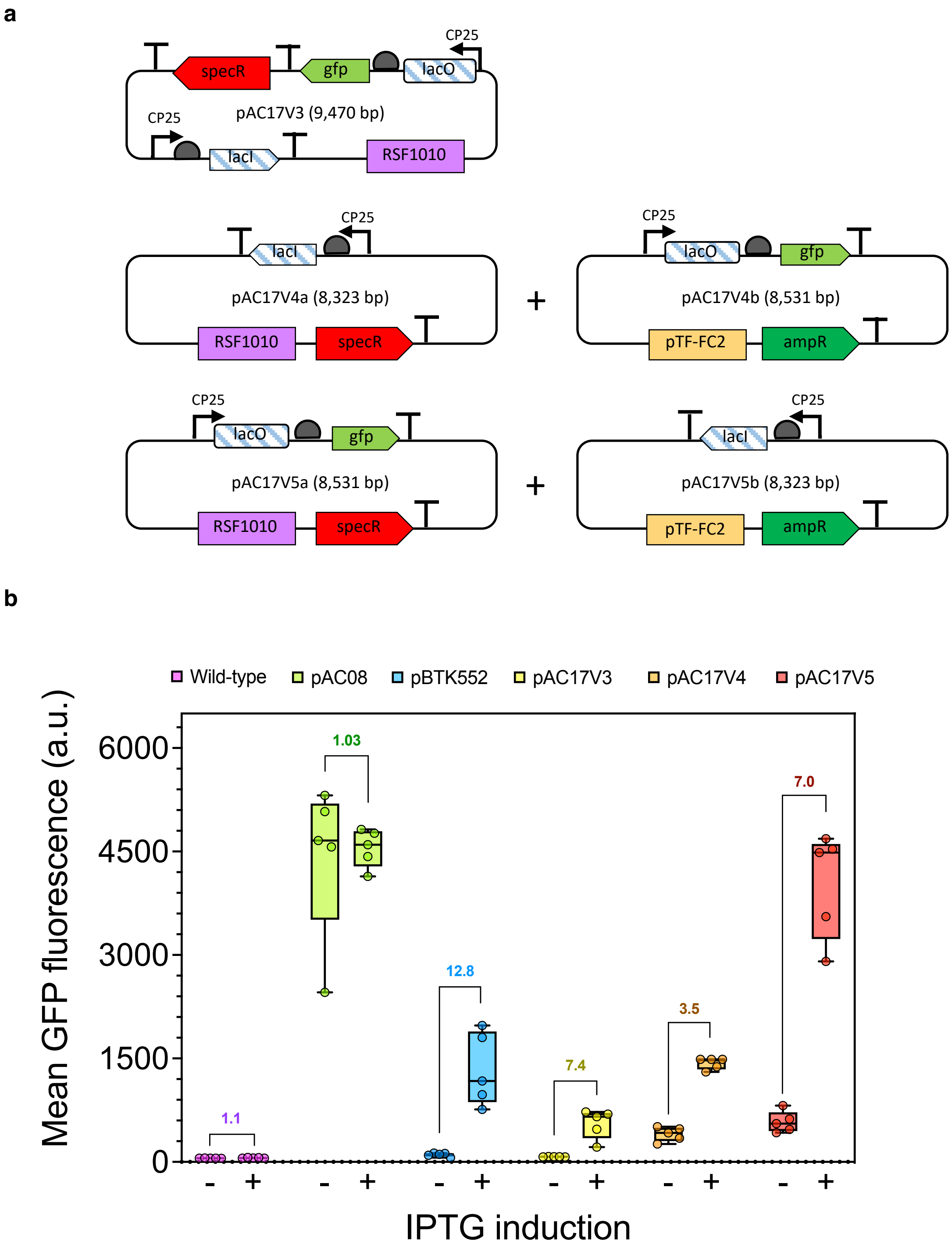


**Supplementary Figure 6. Testing of different IPTG-inducible constructs in *S. alvi*. a** Maps of the IPTG-inducible plasmids built in this study. **b** *S. alvi* cells engineered with our inducible plasmids respond to IPTG exposure *in vitro*. Graph shows box plots representing median value of GFP fluorescence of five biological replicates for each construct tested. Each replicate value is based on the average fluorescence of at least 9,000 *S. alvi* cells measured by flow cytometry, which were grown in liquid with (+) or without (-) IPTG. As a reference, wild-type *S. alvi*, *S. alvi* bearing the pAC08 plasmid constitutively expressing GFP and *S. alvi* carrying the previously built pBTK552 vector (Leonard *et al.*, 2018)[27] were also analyzed. Fold-changes of average fluorescence between uninduced and induced cells are indicated.

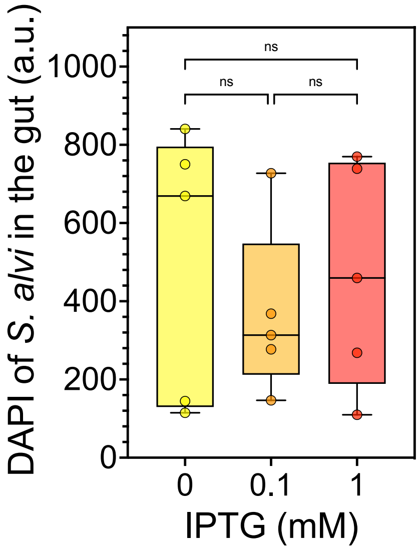


**Supplementary Figure 7. DAPI measurements from gut tissues.** Graph shows box plots representing median value of DAPI fluorescence of *S. alvi* biofilms imaged from the gut of bees fed sugar water supplemented with either 0, 0.1 or 1 mM IPTG. Five bees were analyzed for each condition and fluorescence values were averaged from three distinct sections of each gut. One-way ANOVA test, not significant (ns) with $q$-value > 0.5.

**Fiji macro**

```
dir=getDirectory("Choose Source Dir");
File.makeDirectory(dir+"results/");
File.makeDirectory(dir+"results/GFPmax/");
list=getFileList(dir);
Array.show(list);

function makeCellmask(id){
				print("Creating cell mask from GFP channel for " + id);
				open(dir + id);
				selectWindow(id+" - C=1");
				close();
				selectWindow(id+" - C=0");
				run("Z Project...", "projection=[Max Intensity]");

				run("8-bit");
				run("Duplicate...", "title=["+id+"- Cell mask]");
				run("Duplicate...", "title=["+id+"- Cell peaks]");

				selectWindow(id+"- Cell mask");
				setAutoThreshold("Triangle dark");
				setThreshold(19, 255);
				run("Convert to Mask");

				selectWindow(id+"- Cell peaks");
				run("Duplicate...", "title=["+id+"- Cell signal]");
				saveAs("Tiff", dir+"results/"+id+"- Cell signal");
				wait(500);
				run("Close");

				selectWindow(id+"- Cell peaks");
				run("Median...", "radius=1");
				run("Gaussian Blur...", "sigma=0.20 scaled");
				run("Find Maxima...", "prominence=25 exclude output=[Segmented
Particles]");
				run("Invert");
				wait(500);
				imageCalculator("Subtract create", id+"- Cell mask", id+"- Cell peaks
Segmented");
				saveAs("Tiff", dir+"results/"+id+"- Cell mask");
				wait(500);
				run("Close");
				close(id+"- Cell mask.tif");
				close("\\Others");
				while (nImages>0) {
	selectImage(nImages);
	close();
   }
}

function separatepercell(id){
				print("Creating ROIs for each cell in " + id);
				open(dir +"results/"+ id +"- Cell mask.tif");
```

```
			run("Analyze Particles...", "size=0.8-Infinity circularity=0-1.00 display exclude
clear summarize add");
			run("Clear Results");
			selectWindow("Summary");
			run("Close");
			cells = roiManager("count");
			print("There are " +cells+" cells.");
			wait(1);
			for (c=0; c<cells; c++){
				roiManager("Select", c);
				roiManager("rename", roiManager("index"));
				wait(1);
			}
			roiManager("save", dir+"results/"+id+".zip");
			selectWindow("ROI Manager");
			run("Close");


			for (c=0; c<cells; c++){
				roiManager("reset");
				roiManager("open", dir+"results/"+id+".zip");
				open(dir +"results/"+ id +"- Cell signal.tif");
				wait(50);
				roiManager("Select", c);
				run("Crop");
				run("Measure");
				}
			selectWindow("ROI Manager");
			run("Close");
			selectWindow("Results");
			saveAs("Text", dir+"results/GFPmax/"+id+"_GFPtable.txt");
			wait(50);
			run("Close");
			while (nImages>0) {
				selectImage(nImages);
				close();
			}
			wait(100);
}


			for (l=0; l<list.length; l++){
			print("Starting :"+list[l]);
			if (endsWith(list[l], "/")){
				print("This is a directory");
				}
			else{
			if (endsWith(list[l], ".czi")){
				run("Clear Results");
				id = list[l];
				makeCellmask(id);
				separatepercell(id);
				}
```

```
            }
            wait(1);
            close("\\Others");
            while (nImages>0) {
            selectImage(nImages);
            close();
   }
}
```
